# More or fewer latent variables in the high-dimensional data space? That is the question

**DOI:** 10.1101/2024.11.28.625854

**Authors:** Francesco Edoardo Vaccari, Stefano Diomedi, Edoardo Bettazzi, Matteo Filippini, Marina De Vitis, Kostas Hadjidimitrakis, Patrizia Fattori

## Abstract

Dimensionality reduction is widely used in modern Neuroscience to process massive neural recordings data. Despite the development of complex non-linear techniques, linear algorithms, in particular Principal Component Analysis (PCA), are still the gold standard. However, there is no consensus on how to estimate the optimal number of latent variables to retain. In this study, we addressed this issue by testing different criteria on simulated data. Parallel analysis and cross validation proved to be the best methods, being largely unaffected by the number of units and the amount of noise. Parallel analysis was quite conservative and tended to underestimate the number of dimensions especially in low-noise regimes, whereas in these conditions cross validation provided slightly better estimates. Both criteria consistently estimate the ground truth when 100+ units were available. As an exemplary application to real data, we estimated the dimensionality of the spiking activity in two macaque parietal areas during different phases of a delayed reaching task. We show that different criteria can lead to different trends in the estimated dimensionality. These apparently contrasting results are reconciled when the implicit definition of dimensionality underlying the different criteria is considered. Our findings suggest that the term ‘dimensionality’ needs to be defined carefully and, more importantly, that the most robust criteria for choosing the number of dimensions should be adopted in future works. To help other researchers with the implementation of such an approach on their data, we provide a simple software package, and we present the results of our simulations through a simple Web based app to guide the choice of latent variables in a variety of new studies.

**Key points:**

- Parallel analysis and cross-validation are the most effective criteria for principal components retention, with parallel analysis being slightly more conservative in low-noise conditions, but being more robust with larger noise.
- The size of data matrix as well as the decay rate of the explained variance decreasing curve strongly limit the number of latent components that should be considered.
- When analyzing real spiking data, the estimated dimensionality depends dramatically on the criterion used, leading to apparently different results. However, these differences stem, in large part, from the implicit definitions of ‘dimensionality’ underlying each criterion.
- This study emphasizes the need for careful definition of dimensionality in population spiking activity and suggests the use of parallel analysis and cross-validation methods for future research.

## Introduction

In the last decade, state-space and dimensionality reduction techniques have dominated the data analysis in Neuroscience, especially when studying population spiking activity (Pellegrino et al., 2024; Keemink and Machens, 2019; Paninski and Cunningham, 2018; Pang et al., 2016; Cunningham and Yu, 2014) and have been validated by solid theoretical work (Humphries, 2023; Badre et al., 2021; Jazayeri and Ostojic, 2021; Gallego et al., 2017; Gao et al., 2017). In parallel, computational studies have led to highly sophisticated, non-linear methods mostly based on autoencoder neural networks (for example, LFADS,Pandarinath et al., 2018; pi-VAE, Zhou and Wei, 2020; PSID, Sani et al., 2021; TNDM, Hurwitz et al., 2021; CEBRA,Schneider et al., 2023; d-VAE, Li et al., 2023). Despite their capacity to model neural activity and find latent dynamics related to behavior, the use of such complex tools is not as widespread, mainly because of their complexity that requires a strong background in machine learning and neural networks that is not always the case for most Neuroscientists. Also, depending on the specific application, linear techniques still offer the best trade-off between modelling relevance, simplicity and explainability. Indeed, many works still rely on Principal Component Analysis (PCA), Singular Value Decomposition (SVD) or their derivations with extremely intriguing results (Heller and David, 2022; Keemink and Machens, 2019; Semedo et al., 2019; Williams et al., 2018; Seely et al., 2016; Kobak et al., 2016). For example, very recently Pellegrino and colleagues (2024) developed a compact model (sliceTCA) that is able to demix covariability across neurons, time and trials and provide insights into the neural computations underlying behavior.

An important issue that affects all these methods (both linear and non-linear) is the choice of optimal number of dimensions to retain (i.e., the real dimensionality of a dataset) for visualization and, more importantly, for subsequent analysis. Non-linear models are often cross validated by default to reduce over-fitting, but this is not the case for linear models, thus this issue is even more relevant for them.

To address this issue, here we chose to focus on PCA since it is the simplest, widely used and computationally efficient technique. Normally, an arbitrary number of Principal Components (PCs) (Safaie et al., 2023; Chen et al., 2021; Russo et al., 2018) or the minimum number of PCs required to achieve an arbitrary proportion (often 80% or 90%) of the explained variance (Thura et al., 2022; Maranesi et al., 2019; Elsayed et al., 2016) is retained. More recently, a simple metric calculated starting from the eigenvalues (i.e., the Participation Ratio, PR) has been used (Fortunato et al., 2024; Johnston and Fusi, 2023; Gao et al., 2017; Mazzucato et al., 2016). Other criteria for choosing the best number of PCs have been proposed outperforming the aforementioned ones, but their use in Neuroscience is very limited. Among these, we identified the Kaiser rule (Kaiser, 1960), the parallel analysis (PA; Horn, 1965) and cross-validation as the most promising (see Methods and Discussion for details).

A related study was conducted by Altan and colleagues (2021), where the authors focused on dimensionality estimation with linear and nonlinear algorithms. They found that, as expected, for linear noise-free generated data both types of algorithms performed similarly well, whereas for nonlinear data the linear algorithms performed poorly. However, when noise was intentionally introduced into the system, the nonlinear methods were the most affected, overestimating the real dimensionality to the point that they performed worse than linear ones even on nonlinear data. Thus, the authors concluded that noise can disrupt some methods of dimensionality estimation, but they did not investigate this crucial aspect in depth. Building on these findings, we further investigated the effect of noise on different linear estimation methods related to PCA to provide practical guidelines for population neural recording studies in which noise is unavoidable. Similarly to Altan and colleagues, we here compared different criteria for dimensionality estimation on simulated data. In this case, simulations are essential to have ground-truth data, both in terms of dimensionality and noise amount. Simulations also allow us to manipulate several parameters of the data to derive suggestions about future real applications. In this regard, various scenarios were conceived to provide insights useful for a general-purpose reader, but also tuned to mimic neural activity. Finally, we applied different criteria for dimensionality estimation to real spiking activity to highlight the differences in the results they may generate.

This work is intended as a simple guide for neuroscientists using PCA in choosing the most appropriate number of com-ponents and proposes to collectively adopt the criteria that performed best in our simulations as a new standard. Accordingly, a simple software package is provided to facilitate the adoption of these methods in the community, thus improving results comparability.

## Materials and Methods

The primary scope of this work is to provide guidance for the analysis of neural data. However, the criteria used to estimate the optimal dimensionality are not necessarily derived from the neuroscience literature (in fact, most of them are not), and can therefore be applied to a wide range of datasets. Although, in the following sections, we refer to the columns of the synthetic data matrices as ‘units’ or ‘neurons’ for clarity, they are simply time series generated as linear combinations of a set of latent variables, thus they can be representative of different types of data (e.g., EMG recordings; see Discussion).

Finally, in the following paragraphs, when we refer to ‘the first PCs’ or similar terms, we mean the first PCs sorted according to their explained variance.

### Simulations

To compare the performance of different criteria for choosing the optimal number of latent variables to retain we applied them on a variety of synthetic data. This step was unavoidable to have a ground truth underlying structure knowing a priori the number of components and their temporal dynamics as well as the amount of noise injected into the system. Moreover, by controlling the different parameters of the data generation procedure we have been able to simulate various scenarios in which the different criteria of interest were tested.

The simulation procedure, similar to that used in Altan and colleagues (2021), followed a number of steps: 1) latent variables generation; 2) linear combination to create time series; 3) adding noise; 4) normalization.

#### 1. Latent Variables Generation

The latent variables for generating the data matrix were generated following two alternative procedures to avoid biases: i) random dynamical process; ii) using PCs of real datasets.

i. The first option was using a simple iterative algorithm described by:

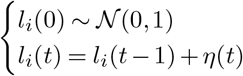

where *l*_*i*_(*t*) is the value of the *i*^*th*^ latent variable at time *t* and *η*(*t*) ∼ 𝒩 (0, 1) is a value drawn from a gaussian distribution with mean equal to 0 and standard distribution equal to 1.
ii. Alternatively, latent variables were directly extracted from real neural data via PCA (second option). Spiking activity recorded from medial parietal area V6A was recorded while a macaque performed an instructed delay reaching task. This data is available online as a part of our recently published dataset https://doi.gin.g-node.org/10.12751/g-node.7q2dbp/ (Diomedi et al., 2024b; for more details about recordings, see Diomedi et al., 2024a). Briefly, neural data was binned at 25ms, averaged across conditions, soft-normalized (see below) and PCs were extracted with PCA. To generate new data, a subset composed of *N* (see below) of these real PCs was randomly selected each time. This second variant of the procedure ensured that the final simulated time series mimicked every aspect of real neural activity.

At this stage, we could vary two parameters, specifically the number of latent variables to be generated (*N*) and the length of each time series (*T*).

The subset of latent variables -obtained either through procedure i) or ii)- to be used for data generation was first z-scored, then orthogonalized via PCA (i.e. running PCA on the matrix of latent variables and retaining for subsequent steps the resulting score matrix) to ensure the complete independence of latent variables. Finally, all latent variables were first z-scored to obtain a set of variables with equal variance (= 1). Each of orthogonalized, z-scored latent variables were then rescaled to obtain a monotonically decreasing variance. This step was motivated by the fact that latent variables (i.e., PCs) in natural, real data show a decrease in variance after PCA. For each *i*^*th*^ latent variable, we computed the scaling factor *SF*_*i*_ drawn from an exponential decay curve as follow:

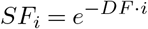

where the decay factor *DF* can be regulated during the simulation.

For each simulation, the scaling factors were then normalized to ensure comparability across different simulations:

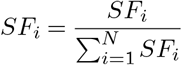

where *N* indicates the number of generated latent variables. During the step of scaling, the parameter *DF* is crucial for regulating the steepness of the decay curve and thus affecting the amount of variance explained by each of the generated latent variables in the final data matrix. Specifically, when *DF* increases, the first latent variables will have a much greater importance than the last ones. For even higher *DF*, the last latent variables will be irrelevant to the point that they will not be detectable in the final matrix and be confused with the noise.

#### 2. Linear Combination to Create Time Series

Once the set of latent variables was properly orthogonalized and scaled, we generated a matrix *W* ∈ ℝ^*M ×N*^ containing *M* random vectors (each containing *N* random values drawn from a uniform distribution in the range [− 0.50.5], followed by normalization of each vector) to generate the activity of *M* synthetic neurons as linear combinations of the generated latent variables:

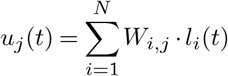

where *u*_*j*_(*t*) is the value representing the activity of the *j*^*th*^ synthetic neuron at time *t*. At the end of this step, the vector *u*_*j*_ was z-scored to obtain the same variance (= 1) for all synthetic units. We stacked together the *u*_*j*_ vectors to generate the *U* ∈ ℝ^*T ×M*^ matrix containing the synthetic neurons noise-free activity.

#### 3. Adding Noise

In this step, a random noise matrix *E* ∈ ℝ^*M ×N*^, whit the same size as the matrix containing the synthetic unit data, was generated by drawing its elements from a Gaussian distribution 𝒩 (0, 1). At this point, each column of both the data matrix and the noise matrix had equal variance (= 1). To control the amount of noise to be added to the data matrix, the noise matrix was scaled by a Noise Factor (*NF*). When *NF* ≈ 0, the amount of noise was insignificant, while for *NF* ≈ 1 the noise had the same variance as the signal (i.e., the generated data matrix), representing approximately the 50% of the total variance in the final matrix. Although such high noise values may seem unphysiological, our application on real data showed even higher values (see Results). Further reasons for this noise range are provided in the Discussion.

Since it is unlikely that all units are equally noisy in real situations, we composed the final matrix *X* as follows:

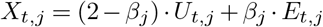

where *U* is the matrix with synthetic unit data, *E* is the matrix of rescaled gaussian noise and *β*_*j*_ is a parameter that controls the amount of noise added to each synthetic unit. Note that we chose the constant value 2 for simplicity, so that when *β*_*j*_ = 1, the ratio of neural activity to noise was 1 : 1 (when *β*_*j*_ = 0.5, the ratio was 1.5 : 0.5, and so on). This approach would be equivalent if we chose 1 − *β*_*j*_ as the weighting factor instead. For a subset of simulations, we set *β*_*j*_ = 1 for all synthetic units, obtaining equally noisy synthetic units (similarly to the noise added in Altan et al., 2021). However, since real neural data often include a mix of more and less noisy units, we set 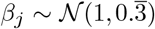 for another subset of simulations, with the additional constraint that *β*_*j*_ remains within the range [02]. To ensure that *X* was completely positive matrix, as is typical in real spiking neural activity, and without affecting the latent variables’ temporal patterns, we performed the transformation:

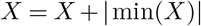

At the end of the procedure, *X* ∈ ℝ^*T ×M*^, with each row representing a time bin and each column a synthetic neuron. Formatting the data as a matrix was necessary to subsequently apply PCA.

### 4. Normalization

Normally, it is good practice to standardize data before PCA (e.g., via z-score normalization), and this was our first option. However, when dealing with real neural data, soft-normalization is preferable because it is a good trade-off between hard normalization (where each unit will have the same weight in determining PCs, regardless of spiking activity variance) and no normalization (which is biased towards the more active units in the population). Since soft-normalization is widely used in Neuroscience applications involving state-space methods and dimensionality reduction (Jiang et al., 2020; Elsayed et al., 2016; Churchland et al., 2012), we used soft-normalization instead of z-scoring in a subset of simulations before applying PCA and criteria for selecting the PCs. Soft-normalization was performed as follows:

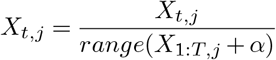

where *X*_1:*T*,*j*_ is the noisy data of the *j*^*th*^ synthetic unit over *T* time points, and *α* is a constant set equal to the mean of *X* (i.e., the mean activity of the simulated population). After soft-normalization, we subtracted from each *j*^*th*^ column its mean.

In summary, we designed a procedure to build synthetic data matrices of size [*T, M*] containing *M* synthetic unit responses obtained by linearly combining *N* latent variables from either a random dynamical process or real neural dynamics. Other adjustable parameters included the steepness of the latent variable variance decreasing curve (*DF*), the noise level (*NF*), the noise distribution across the population of synthetic units (*β*), the type of matrix normalization (z-scoring or soft-normalization).

Importantly, at the end of the simulation process, we computed the decay rate *τ* of the explained variance associated with the PCs of the final matrix *X*. Similarly to the scaling factor *SF* equation, we expected the explained variance to decrease following:

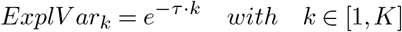

where *ExplV ar*_*k*_ is the explained variance associated to the *k*^*th*^ PC and *K* is the total number of PCs (normally equal to *M*, the number of synthetic units). Note that *τ* (calculated a posteriori from the final data matrix) is related to *DF* (set for a specific simulation), but it is impossible to know *τ* exact value a priori due to randomness. Nevertheless, *τ* has an impact on then number of latent variables that may be estimated in the data (see Fig. 5 in the Results).

### Criteria for dimensionality estimation

In our study, we compared multiple criteria to estimate the optimal number of PCs:

**Table 1.**
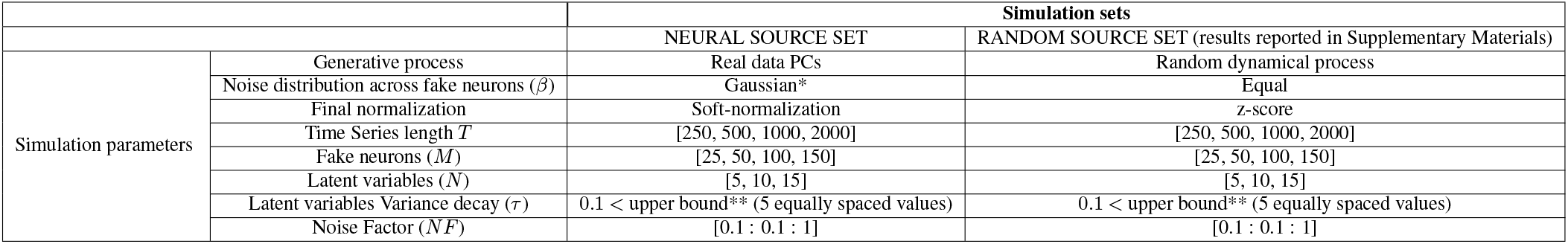
Simulation parameter settings for the two simulation sets RANDOM SOURCE and NEURAL SOURCE. The remaining values are defined in the Methods section. * See Methods for Gaussian distribution values and constraints. ** the upper bound was calculated so that the latent variable with the lowest explained variance still accounted for more variance than the first PC consisting of pure noise.

|  |  | Simulation sets |  |
| --- | --- | --- | --- |
|  |  | NEURAL SOURCE SET | RANDOM SOURCE SET (results reported in Supplementary Materials) |
| Simulation parameters | Generative process | Real data PCs | Random dynamical process |
| | Noise distribution across fake neurons ($\beta$) | Gaussian* | Equal |
|  | Final normalization | Soft-normalization | z-score |
| | Time Series length $T$ | [250, 500, 1000, 2000] | [250, 500, 1000, 2000] |
| | Fake neurons ($M$) | [25, 50, 100, 150] | [25, 50, 100, 150] |
| | Latent variables ($N$) | [5, 10, 15] | [5, 10, 15] |
| | Latent variables Variance decay ($\tau$) | 0.1 < upper bound** (5 equally spaced values) | 0.1 < upper bound** (5 equally spaced values) |
| | Noise Factor ($NF$) | [0.1 : 0.1 : 1] | [0.1 : 0.1 : 1] |

#### i) Explained Variance Threshold and Participation Ratio (PR)

First, since they are commonly used, we also implemented two explained variance thresholds, one at 90% and another at 80%. Additionally, we used the Participation Ratio (PR) calculated as follows (Gao et al., 2017; Mazzucato et al., 2016):

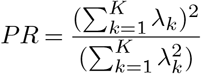

where *λ*_*k*_ is the eigenvalue associated to the *k*^*th*^ PC, and *K* is the total number of PCs (normally equal to *M*, the number of synthetic units). A *PR* = 1 indicates that all the variance is explained by the first PC (thus *λ*_*k*_ = 0 for *k >* 1), while a *PR* = *K* = *M* implies that variance is equally distributed across all PCs (thus *λ*_1_ = *λ*_2_ = … = *λ*_*k*_). Values within these lower and upper bounds indicate the concentration of the variance in the first PCs, providing a direct estimation of the number of PCs to retain. This criterion generally behaves similarly to a 75 − 77% variance threshold (Gao et al., 2017; see also our Results).

#### ii) Kaiser rule

In addition, we applied the Kaiser rule, which suggests to retain all PCs with eigenvalues greater than one (Dinno, 2014; Kaiser, 1960). The reasoning behind this criterion is that eigenvalues greater than 1 correspond to PCs which explain more variance than each of the original variables. However, the relationship between eigenvalues and explained variance is straight-forward when data are standardized before applying PCA (i.e., z-scored), but it is not when data have been soft-normalized (since different original variables can explain more variance than others). Since we used z-scored, but also soft-normalized data, we adapted the Kaiser rule to retain all the PCs explaining more variance of *X* than the average variance explained by each individual column (i.e., unit) in *X*.

#### iii) Parallel analysis

Parallel analysis is a derivation of the Kaiser rule (Dinno, 2014, 2009; Hayton et al., 2004) that instead of using a single, fixed threshold for the eigenvalues (= 1), it uses values derived from a random null distribution. We generated a surrogate matrix by reshuffling the elements of each column (synthetic unit) of the matrix *X*, thus preserving individual columns statistics, but disrupting any significant correlation due to the latent structure except for the spurious ones. We sorted in descending order the eigenvalues of the covariance matrix of these surrogate data. This procedure was normally repeated 250 times when not otherwise specified (100 and 1000 iterations were also tested in a subset of simulations with negligible differences) obtaining a null distribution of eigenvalues from surrogate matrices. Finally, a PC was retained if its associated eigenvalue was higher than the 95^*th*^ percentile of the eigenvalue null distribution.

#### iv) Cross-Validation with imputation

The last method we selected for choosing the optimal number of PCs is a k-fold cross-validation scheme. This technique is commonly used in machine learning and also for estimating the number of latent variables for most of the non-linear dimensionality reduction algorithms. However, it is rarely applied for PCA due to the complexity of cross-validation in dimensionality reduction, where the model must be able to extract latent dynamics from the training data robust enough to generalize to both new time points and new units. Thus, differently from the majority of models (for example, regressions), in the framework of dimensionality reduction there is the need not only to exclude from the training set data along the rows of the *X* matrix (i.e., time points), but also data along the columns (i.e., synthetic units)(Bro et al., 2008; Diana and Tommasi, 2002; Wold, 1978). This procedure has the advantage of avoiding bootstrapping but has the drawback of producing a training matrix with missing values. Since the algorithms for PCA might become non-efficient and complex when dealing with data with missing values, we implemented an iterative algorithm based on imputation (similarly to what suggested by Kiers, 1997). In our approach, randomly selected elements of *X* are removed and set apart as test set. These elements are replaced by 0s (i.e., the mean of *X*, since the matrix must be centered subtracting the mean), allowing us to use classic PCA algorithms without missing values. Then, the selected elements (but not all the other elements constituting the training set) are updated at the end of each iteration until convergence. The following pseudo-code outlines our iterative cross-validation procedure:

1. FOR each *CV* in [1 *NumberOf CV f olds*]:
  a. Indices = random selection of elements from the data matrix *X*
  b. FOR each *d* in [1 *MaxDimensions*]:
    i. Initialize the matrix *Z*_*k*_ = *X*
    ii. Set *Z*_*k*_(*Indices*) = 0
    iii. Initialize the matrix *Z*_*k−*1_ with random values (same shape as *Z*_*k*_)
    iv. FOR each *k* in [1 *MaxIterations*]:
      - Perform PCA on *Z*_*k*_ (scores and coefficient vectors as output)
      - Update the *Z*_*k*_(*Indices*) by projecting the first d PCA scores onto the first d PCA coefficient vec-tors
      - 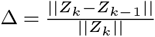
      - Update *Z*_*k−*1_ = *Z*_*k*_
      - *k* = *k* + 1
      - IF Δ *< tol* (reached convergence): STOP
    v. Calculate *R*^2^(*CV, d*) between *Z*_*k*_(*Indices*) and *X*_*k*_(*Indices*)
2. Average *R*^2^(1 : *NumberOf CV f olds, d*) values for each *d* across all cross-validations.
3. Optimal *d* = *argmaxR*^2^(*d*)

Note that we considered that the algorithm converged when the relative change Δ in *Z* between consecutive iterations *k* and *k* + 1:

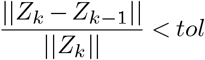

where *tol* is an arbitrary low threshold (tolerance). Since the algorithm updates only the test elements of the matrix, only these will be changing from an iteration and the next one.

Normally, when cross validation is used to tune one hyperparameter of an algorithm, the performance is expected to peak at the optimal value for that hyperparameter and to decrease as values move away from this point. In our case, we considered the number of PCs as an hyperparameter, thus we expected that the *R*^2^ of the test elements reconstruction would increase with the number of PCs peaking in correspondence with the optimal number of PCs. Beyond this point, adding more components leads to a decrease in the *R*^2^, meaning that additional PCs capture more noise than significant information. For our implementation, we used a 5− fold cross validation scheme, setting *tol* value to 10^*−*4^ and the maximum number of iterations to 100. Tests with a 10− fold cross − validation scheme and *tol* equal to 10^*−*6^ produced similar results (data not shown). Although this method is computationally expensive compared to other techniques, it runs within a couple of minutes on a common laptop (performance depends on the matrix size).

### Metrics for criteria comparison

We calculated three metrics to evaluate the performance of each criterion, considering three objectives that an optimal method should simultaneously aim to achieve: i) estimate the correct number of latent variables underlying the data, ii) provide a 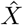 denoised ma-trix (reconstructed using only the first PCs according to the method selection,) which retains as much information as possible related to the noise-free data matrix *u*, iii) estimate the correct amount of noise (i.e., uncorrelated variations across different units; see Discussion).

Specifically, we calculated: i) the dimension error, calculated as the difference between the estimated number of PCs and the number of latent variables *N* used to generate *X*; ii) the information about the latent variable retained in the reconstructed matrix, calculated as the variance in the reconstructed 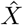 that is explained by *u* (i.e., the matrix obtained by linear combinations of the generated latent variables); iii) the estimated noise error, calculated as the difference between the proportion of variance of in *X* unexplained by the generated latent variables *l* (ground-truth noise amount) and the proportion of variance of *X* unexplained by the selected PCs (noise estimated by the used criterion).

### Neural data pre-processing

To further validate the applicability of these methods to real world situations, we analyzed spiking data available online (Diomedi et al., 2024b,a) that were recorded from parietal cortex of macaques during an instructed delay reaching task towards visual targets located at various distances and directions in the peripersonal space. We included all neurons recorded across for 10 correct trials / per target were retained. Briefly, we aligned spikes were aligned toon the task events of interest, and we activity was binned the activity into 50ms non-overlapping windows. Neural activity was then averaged for each target, soft-normalized (*E* = 5 *sp/s*) and mean-centered. A We obtained a matrix *X* (size [9 · *T, M*]) was obtained for each task phase of interest., focusing on Specifically, we considered the two key phases: when the animal had its hand in the starting position and waited for the visual target to switch on (FREE; from trial beginning to 400ms after) and during arm movement execution (MOVE; from home button release to 400ms after).

## Results

An ideal method for choosing the optimal number of latent components to retain should be able to estimate not only the number of relevant dimensions, but also the amount of noise (here defined as the uncorrelated variance or “fluctuations unique to each neuron” according to Humphries, 2023) independently. Moreover, the dimensionality and the noise amount estimates should be invariant with respect to the number of units.

For testing different criteria, we first implemented a procedure to generate matrices with a known underlying structure composed of a defined number of latent variables. This process was designed to be flexible, allowing us to simulate various scenarios. Specifically, the parameters that could be varied were: final matrix size [*T, M*] where *T* represents the number of observations, i.e. time bins, and *M* the number of variables, i.e. synthetic neurons), the number of latent variables N and the way they were generated (either with a random dynamical process or starting from real neural data), the steepness of the explained variance decreasing curve *DF* which affects the shape of the eigenspectrum, the amount of noise *NF*, the distribution of noise across the population of synthetic units *β*, and the type of matrix normalization (z-scoring or soft-normalization).

Results are organized in 3 different paragraphs: 1) simulations to compare different criteria and identify the most robust methods for estimating the dimensionality of a dataset; 2) simulations for estimating the upper bound for the number of identifiable components in real situations i.e. when the number of latent variables is unknown; 3) application of these methods to real spiking data.

### Criteria comparison

In this section, we aim to compare the performance of various PCs selection criteria on matrices that contain a predefined number of significant latent variables (thus the dimensionality of the matrix is known). Indeed, during the data simulation procedure, the final matrices were obtained by adding a ‘data matrix’ derived from known latent variables and a ‘noise matrix’ (uncorrelated and randomly generated). Importantly, the steepness of the explained variance decreasing curve of the latent variables was constrained varying*DF* to ensure that the *N* ^*th*^ latent variable (i.e., the last one when ordered for explained variance) accounted for more variance of *X* than the *N* + 1^*th*^ component (i.e., the first component containing pure noise). In this way, the decreasing variance explained by the latent variables was never lower than the highest noise component, ensuring that the number of detectable PCs was effectively equal to *N* .

We focused on a subset of parameters to simulate data that could mimic real neural activity, such as generating latent variables from previously recorded spiking data, setting the noise to be normally distributed over the synthetic units, and soft-normalizing the final matrix (NEURAL SOURCE SET). To generalize findings, we also ran another set of simulations generating latent variables via a random dynamical process, setting equal noise across all observed variables, and z-scoring the final matrix (RANDOM SOURCE SET), useful for datasets of other types, such as EMG (see Discussion). The results from the RANDOM SOURCE SET are reported in the Supplementary Materials and led to very similar findings to the NEURAL SOURCE SET.

The other simulation parameters were set according to the values in Table 1. We ran 5 simulations for each combination of the parameters, generating new data every time. The performance metrics we calculated were: i) the dimension error as the difference between the estimated dimensionality and predefined number of latent variables *N* ; ii) the information about the latent structure retained in the denoised matrix expressed as a fraction of variance of the denoised matrix; iii) the estimated noise error (see Methods for further details). Note that the last two metrics are expected to be inversely correlated, as a method that can reliably discriminate noise (thus providing a low estimated noise error) will likely also be able to retain only the proper information in the denoised matrix.

To report the results, we computed the mean absolute error (MAE) of the dimension error and the noise estimation (accounting for their positive and negative values) as well as the mean for the information retained over all simulations. In particular, these values may not give a complete picture of how the different methods perform in different situations, so in the following paragraphs we will report the results in the figures as a function of noise.

Fig. 1 reports the three-performance metrics calculated over approx. 12000 simulations of the NEURAL SOURCE SET (see Fig. S1 for RANDOM SOURCE SET results). Overall, the 90% and 80% variance thresholds performed well only in a restricted range of noise around the chosen threshold. Indeed, for lower noise levels, they tended to underestimate the dimensionality of the data, excluding relevant information, whereas for higher noise levels they consistently overestimated the dimensionality of the data (Fig. 1A, grey and green lines), incorporating an increasing amount of noise in the ‘denoised’ matrix. Accordingly, the mean absolute error (MAE) for dimensionality estimation was quite high, with values of 19 and 8 for the 90% and 80% thresholds respectively. This pattern (i.e., overestimating noise under the threshold and underestimating above it) is evident in the inverse U-shape of the information retained in the denoised matrix (calculated as variance explained by the known latent structure), which showed a peak at the 10% and 20% noise levels, respectively and decreased at lower and higher values (Fig. 1B, grey and green lines; mean = 74% [78%] for the 90% [80%] thresholds respectively, results averaged across noise levels). By choosing the threshold a priori, these criteria therefore absolutely preclude the possibility of estimating the noise independently of the PCs count, as reflected in a linearly decreasing error in the noise estimation (MAE = 18% [13%] for 90% [80%] thresholds respectively; Fig. 1C, grey and green lines). The Participation Ratio (PR) is related to the variance threshold criteria, but it is based on the calculation of how much the variance is concentrated in the first PCs, thus being more flexible than a fixed threshold. It showed a better performance overall, with fewer PCs retained at higher noise levels (MAE = 3; Fig. 1A, purple line), and a good reconstruction for wider range of noise (peak around 20% – 25%, as expected; mean =83%; Fig. 1B, purple line). How-ever, also PR tended to incorporate more PCs as the noise increased, gradually shifting from dimensionality underestimation to overestimation with a optimal point around 25% noise (Fig. 1A). In addition, it did not provide a good noise estimation, as shown by the linear decreasing error curve (MAE = 8%; Fig. 1C, purple line). The Kaiser Rule performed even better, providing a good estimate of the data dimensionality less sensitive to noise regime (MAE = 2; Fig. 1A, yellow line), a remarkably good data reconstruction (mean = 85%; Fig. 1B, yellow line), and a noise estimate closer to the ground truth (MAE = 5%, Fig. 1C, yellow line). However, also this method tended to include more PCs and to provide less accurate noise estimates as noise increased beyond 25% (Fig. 1A,Cyellow lines). Finally, the cross-validation and the parallel analysis proved to be the most efficient criteria. Both reliably estimated the number of PCs without including more dimensions at high noise levels (MAE = 2 [3] for CV [PA], respectively; Fig. 1A, dark blue and red lines respectively). The data reconstruction was consistently high across the different noise levels (mean = 87 [86%] for CV [PA], respectively; Fig. 1B, dark blue and red lines respectively). Moreover, their capacity to estimate the noise was extremely accurate and consistent across the whole noise range (MAE = 1.4 [3%] for CV [PA], respectively; Fig. 1C, dark blue and red lines respectively). Interestingly, the cross-validation slightly outperformed PA at lower noise levels (0 *−* 25%) incorporating more PCs and providing better data reconstruction, whereas for higher noise regimes (35 *−* 50%) the trend was inverted.

**Fig. 1.**
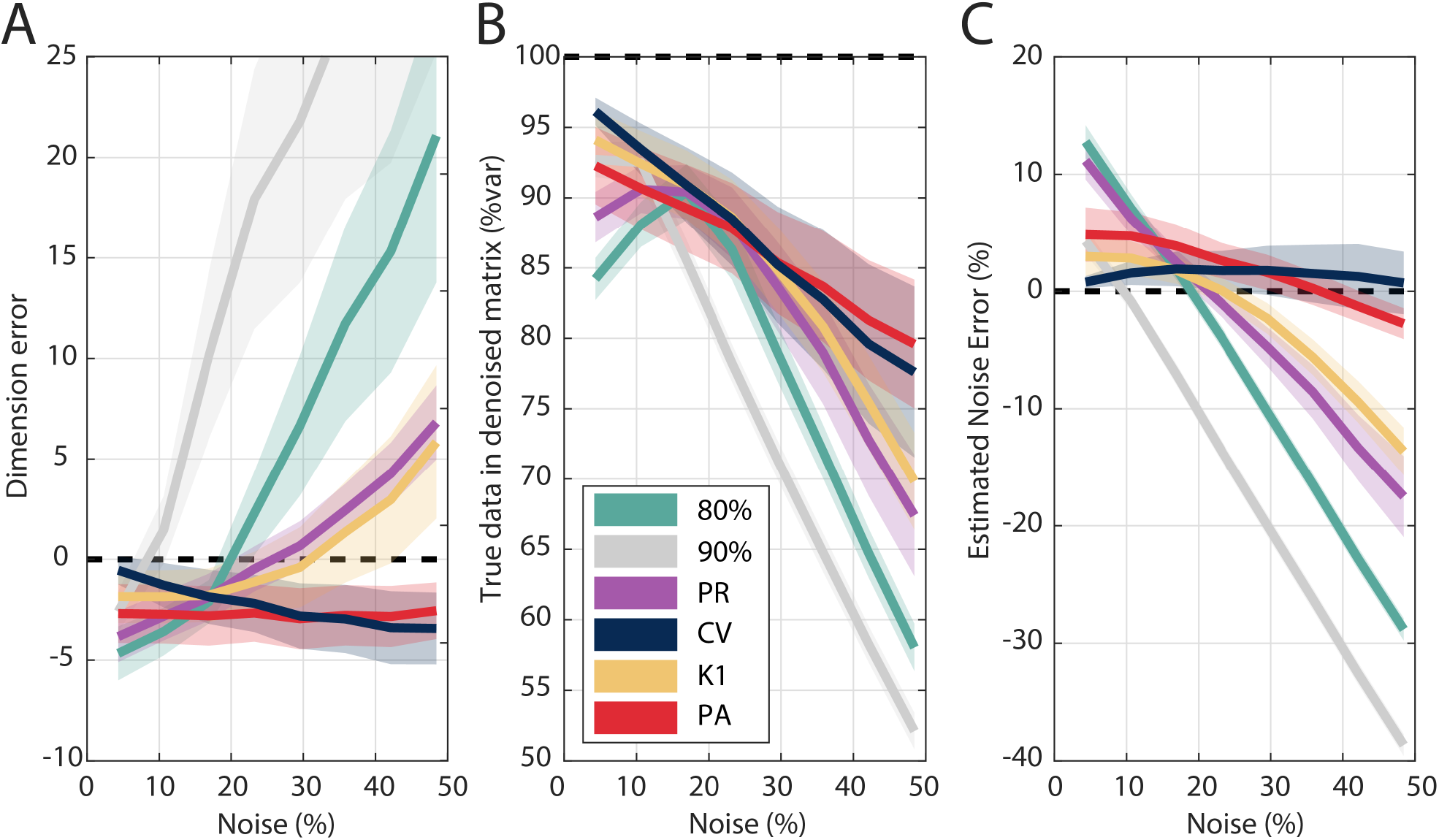
Performance of different criteria for selecting the optimal number of Principal Component tested on simulated data (NEURAL SOURCE SET) as a function of noise (simulation parameters were set according to Table 1). The metrics calculated were: the dimension error, i.e. the difference between the estimated number of PCs and the number of latent variables *N* used to generate the data (A); the information about the latent variables retained in the denoised matrix (B); the estimated noise error (C). Thick lines represent the mean values over simulations. Shaded areas represent mean ± half standard deviation over simulation. Criteria abbreviations: ‘90%’, 90% explained variance threshold; ‘80%’, 80% explained variance threshold; ‘PR’, Participation Ratio; ‘CV’, cross-validated PCA; ‘K1’, Kaiser Rule; ‘PA’, parallel analysis.

Note that while some methods achieved comparable absolute performance scores (e.g., dimension error for PR and PA or information retention for Kaiser Rule and CV), their performance varied significantly across different noise regimes (see also below). We then wanted to compare the performance of the different criteria not only as a function of the noise, but also as a function of the number of latent variables used to generate the data *N*, the number of variables (synthetic units) in the matrix *M* (i.e., columns) and the length of the times series *T* . An optimal method should provide estimates invariant to all these parameters. Figures 2 - 4 report the average results obtained grouping the simulations of the NEURAL SOURCE SET (same as Fig. 1) according to different values of *N*, *M* and *T* (Fig. S2-S4 reports the results for the RANDOM SOURCE SET).

**Fig. 2.**
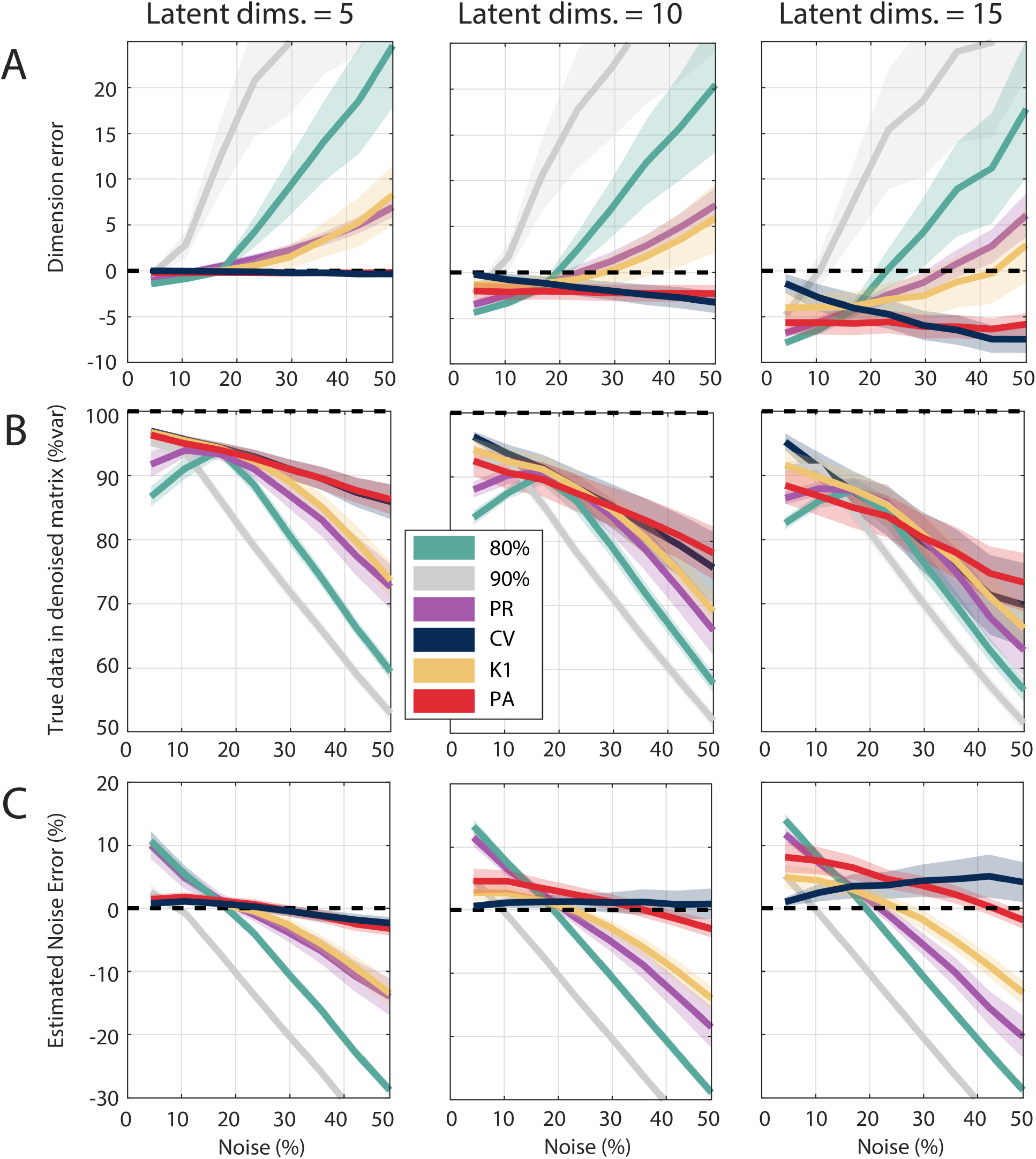
Performance of different criteria tested on simulated data as a function of noise, with results grouped according to the number of latent variables used to generate data matrices (N = 5, 10 or 15). Other simulation parameters were set according to Table 1 (NEURAL SOURCE SET). Other conventions such as in Fig. 1.

When the number of latent variables used to generate the data increased (*N* = [5, 10, 15]), the matrix latent structure became proportionally more complex, resulting in a gradual decrease in the performance for all the criteria. However, the different criteria trends and the relative performance ranking remained mostly unaltered (Fig. 2A-C). PA (but also CV) was the most affected as it is the most conservative method and tended to underestimate dimensionality and overestimate noise as *N* increased. The results obtained for the RANDOM SOURCE SET were in line with these findings (Fig. S2). Note that these results may have been influenced by the choice of DF, which, although it had an upper bound (empirically calculated) to allow all latent variables used in the data generation to be detectable, may not be accurate for larger *N* values, resulting 541 in matrices that have a slightly lower true dimensionality than the desired. When grouping the simulations according to the number of synthetic units (*M* = [25, 50, 100, 150], we observed several major effects. Overall, more synthetic units enhanced sensitivity to noise in estimating the dimensionality (see steeper dimension error vs. noise curves in Fig. 3A) for all criteria except CV and PA. This effect was particularly strong in the variance thresholds methods (Fig. 3A, grey and green lines) and, though present, less pronounced in Kaiser Rule and PR (Fig. 3A, yellow and purple lines). Two factors may explain this phenomenon: first, more components are needed to explain the same amount of variance as the number of units increases (particularly for the hard thresholds and PR), and second, a wider range of eigenvalues is found when the ratio 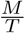 increases (Braeken and van Assen, 2017), leading to more values greater than 1 and thus being included by the Kaiser Rule method. On the contrary, PA and CV were able to leverage the higher amount of data available to estimate more reliably the number of underlying dimensions going from a stable negative dimension error due to its con-servative behavior (Fig. 3 Aleft panels, red and blue lines) to an optimal dimensionality estimation with 100 or more units (Fig. 3 Aright panels, red and blue lines). We want to underline here the robustness of CV and PA, minimally varying their estimates even when *M* is increased sixfold. Regarding information retention in the denoised matrix, either PA and CV leveraged the larger data matrix to perform better (see the gradual upwards shift of red and dark blue lines in Fig. 3B as the number of units increases). Additionally, again PA and CV approached the ground truth in estimating the noise amount, as more units are available (Fig. 3C, red and dark blue lines). Interestingly, the variability for all criteria estimates, but especially for PA and CV was significantly reduced when 100 − 150 units were used (see blue shaded areas in Fig. 3C). This support the use of CV when more data is available. Once more, RANDOM SOURCE SET results (Fig. S3) confirmed this trend. Finally, when simulations were grouped according to the time series length (*T* = [250, 500, 1000, 2000]), we observed negligible effects, except for the Kaiser Rule. In fact, with longer time series, spurious correlations are less likely to occur, leading to a lower sensitivity to noise in dimensionality estimation (see flatter yellow line in Fig. 4A), as well as improved ability to retain the information and estimate noise amount at high noise regimes (Fig. 4B-C, yellow line). This held true also for RANDOM SOURCE SET results (Fig. S4).

**Fig. 3.**
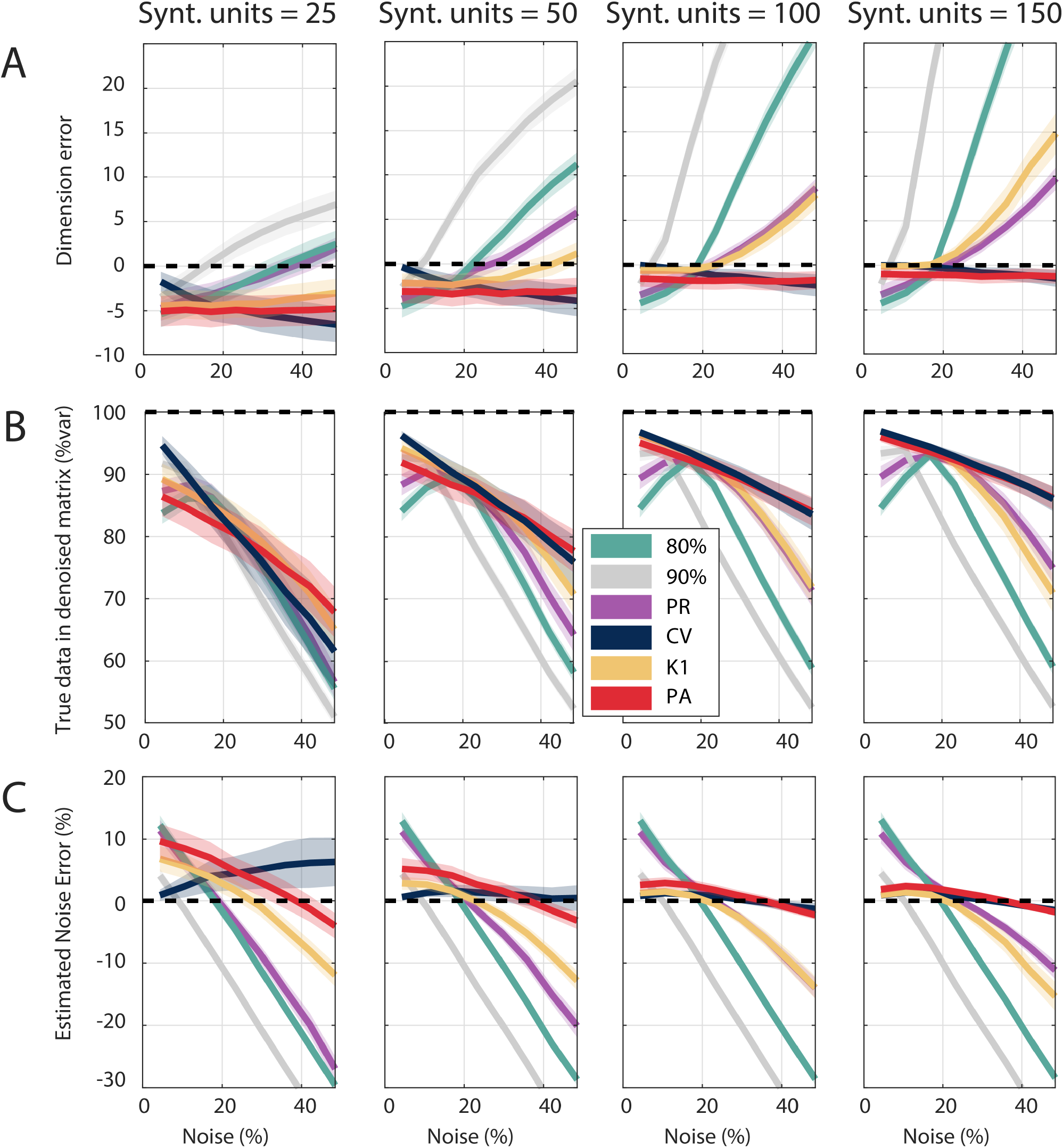
Performance of different criteria tested on simulated data as a function of noise. In this case, results are reported grouping simulations according to the number of synthetic units (M = 25, 50, 100 or 150). Other simulation parameters were set according to Table 1 (NEURAL SOURCE SET). Other conventions such as in Fig. 1.

**Fig. 4.**
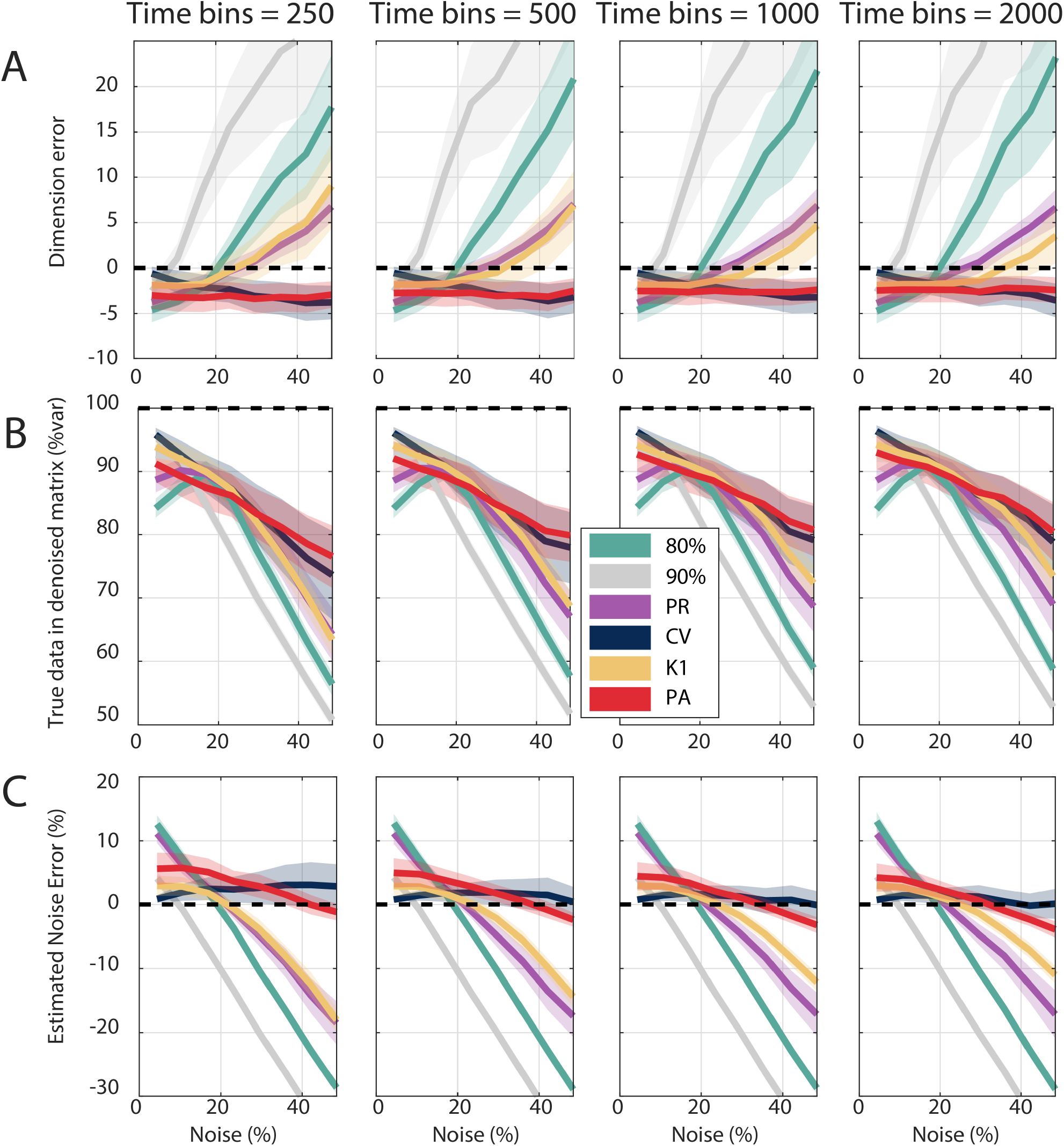
Performance of different criteria tested on simulated data as a function of noise. In this case, results are reported grouping simulations according to the length of the time series (T = 250, 500, 1000 or 2000). Other simulation parameters were set according to Table 1 (NEURAL SOURCE SET). Other conventions such as in Fig. 1.

**Fig. 5.**
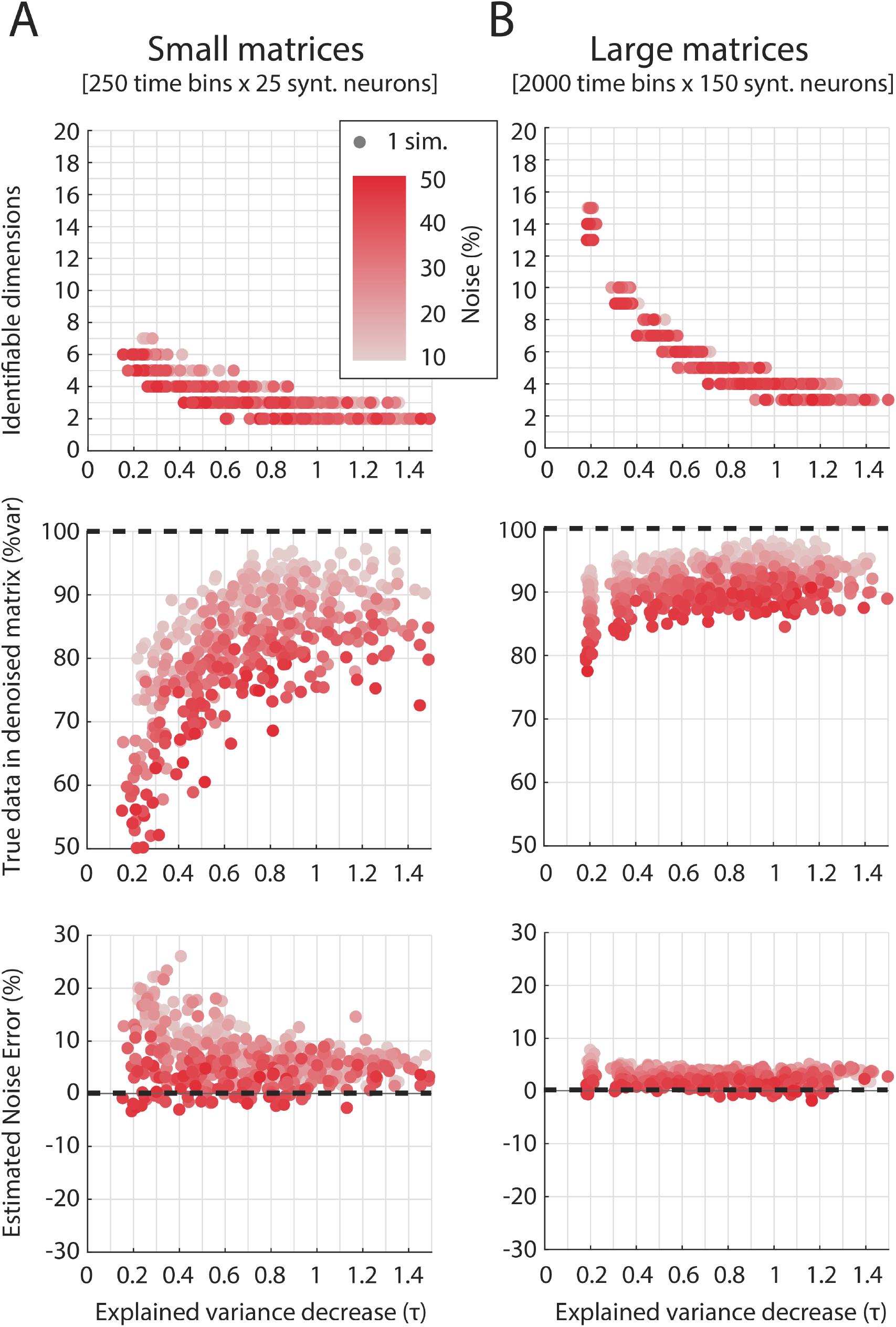
Estimated dimensions, latent variable variance in the denoised matrix and estimated noise for PA in simulations with a high number of latent variables compared to population size. A) 250-timepoints long time series for 25 synthetic units using 20 latent variables were generated and analyzed. B), 2000-timepoints long time series for 150 synthetic units using 80 latent variables. Other parameters were set as NEURAL SOURCE SET in Table 1. Each dot represents a simulation. Color indicates the percentage of noise.

### Simulations for estimating the range of identifiable components

In order to provide some guidelines for future applications, we ran a set of simulations in which data were generated using a very high number of latent variables *N* (*N* = 20 in simulations with 25 synthetic units and *N* = 80 for 150 units; we also set *N* = 40 for 50 units and *N* = 60 for 100 units for other simulations not shown here, but results are available online; see ‘Code availability’). Importantly, in all these cases DFwas not constrained as previously done. This implied that in all simulations, although the latent structure was extremely complex, only a small subset of PCs (with respect to the known *N*) could be reliably estimated after noise was added. The idea behind this analysis was to establish an upper bound for real datasets in which matrix dimensions are known and *τ* can be estimated directly from the real PCs explained variance, but the maximal number of PCs to retain could in principle equal the number of units (i.e., variables in the data matrix). We decided here to focus on PA, as it performed the best across a variety of different scenarios presented above (also when the number of units was limited) and provided results with a very low degree of uncertainty (for results obtained with other criteria, see ‘Code availability’).Fig. 5 shows the results for two example sets of simulations (approx. 500 simulations each set). The left panel shows results for small [250, 25] ([time series length, synthetic units]) matrices, generated using 20 latent variables, while the right panels show results for larger matrices ([2000, 150]) containing 80 latent variables. In Fig. 5, the identified dimensions, the information retained, and the estimated noise error are reported as a function of *τ* . As already mentioned, when *τ* increased, the explainable variance was more concentrated in the first components (Fig. 5A)), allowing fewer latent variables to explain a significant amount of variance and retaining more information in the denoised matrix without incorporating much noise (Fig. 5B). Accordingly, PA showed a monotonically decreasing trend in the number of identifiable dimensions, whereas the variance in the denoised matrix explained by the noise-free data matrix increased. Furthermore, in small matrices (Fig. 5B left), spurious correlations led to fewer significant PCs than in larger matrices (Fig. 5B right). Given a physiological range of *τ* which is normally 0.5 − 0.8 (Qi et al., 2022; Jiang et al., 2020; Kobak et al., 2016), the maximum number of dimensions that can be reliably estimated is 2 − 5 for small matrices and 4 − 8 for larger ones. A few more dimensions could be retained, given that PA is quite conservative and it tends to underestimate the dimensionality, but with the risk to incorporate components with a low signal-to-noise ratio. These findings can have important consequences when interpreting the latent variables in real datasets (see Discussion).

### The dimensionality of neural data depends on the criterion chosen

To show the impact of criterion on dimensionality analysis outcomes, we applied some of the criteria for dimensionality estimation to neural spiking activity from parietal cortex recorded in our laboratory. We analyzed a part of a publicly available dataset (Diomedi et al., 2024a) in which two macaques performed a foveated reaching task that involved a first resting phase in which animals were required simply to hold the arm in the starting position (FREE), a movement planning phase and then the arm reaching execution (MOVE). During the planning phase, monkeys had to maintain fixation at one out of 9 possible targets (placed along 3 directions and 3 depths) that had lit up green, waiting for the GO signal (target color switched red). To investigate the temporal evolution of the neural dimensionality across the task, we compared activity between FREE and MOVE phases, using PR and the PA criteria. PR has been used previously on data recorded from the gustatory cortex of rats and showed that the dimensionality of neural population activity decreases in the presence of a stimulus (Mazzucato et al., 2016). In our case, during MOVE the visual stimulus (absent in FREE) was present and being fixated. Even more importantly, the execution of the arm movement is known to strongly affect parietal activity (Vaccari et al., 2024; Hadjidimitrakis et al., 2019; Cui and Andersen, 2011; Mulliken et al., 2008; Buneo and Andersen, 2006), thus it is likely to affect latent data structure. Among the dimensionality criteria, we compared PR with PA because, based on our simulations (see ‘Criteria comparison’), it performs best with small matrices (in this case, 72 rows). We here focused on Monkey F data (112 and 88 units recorded across 10 correct trials per target from 2 posterior parietal areas V6A and PEc, respectively). To control population size biases, we performed bootstrapping over neurons (1000 repetitions), repeating the analysis each time.

Before performing the dimensionality assessment with PR and PA, we calculated the *τ* across the bootstrapped matrices (sizes [72, 112] for V6A and [72, 88] for PEc). For both areas, *τ* increased significantly from FREE ([0.30, 0.44] for V6A and PEc respectively) to MOVE ([0.41, 0.71] Wilcoxon test, *p <* 0.01; Fig. 6A), indicating that the variance was more concentrated in the first PCs (or, in other words, a few PCs explained a relevant portion of the variance, while the variance explained by the others decrease rapidly).

**Fig. 6.**
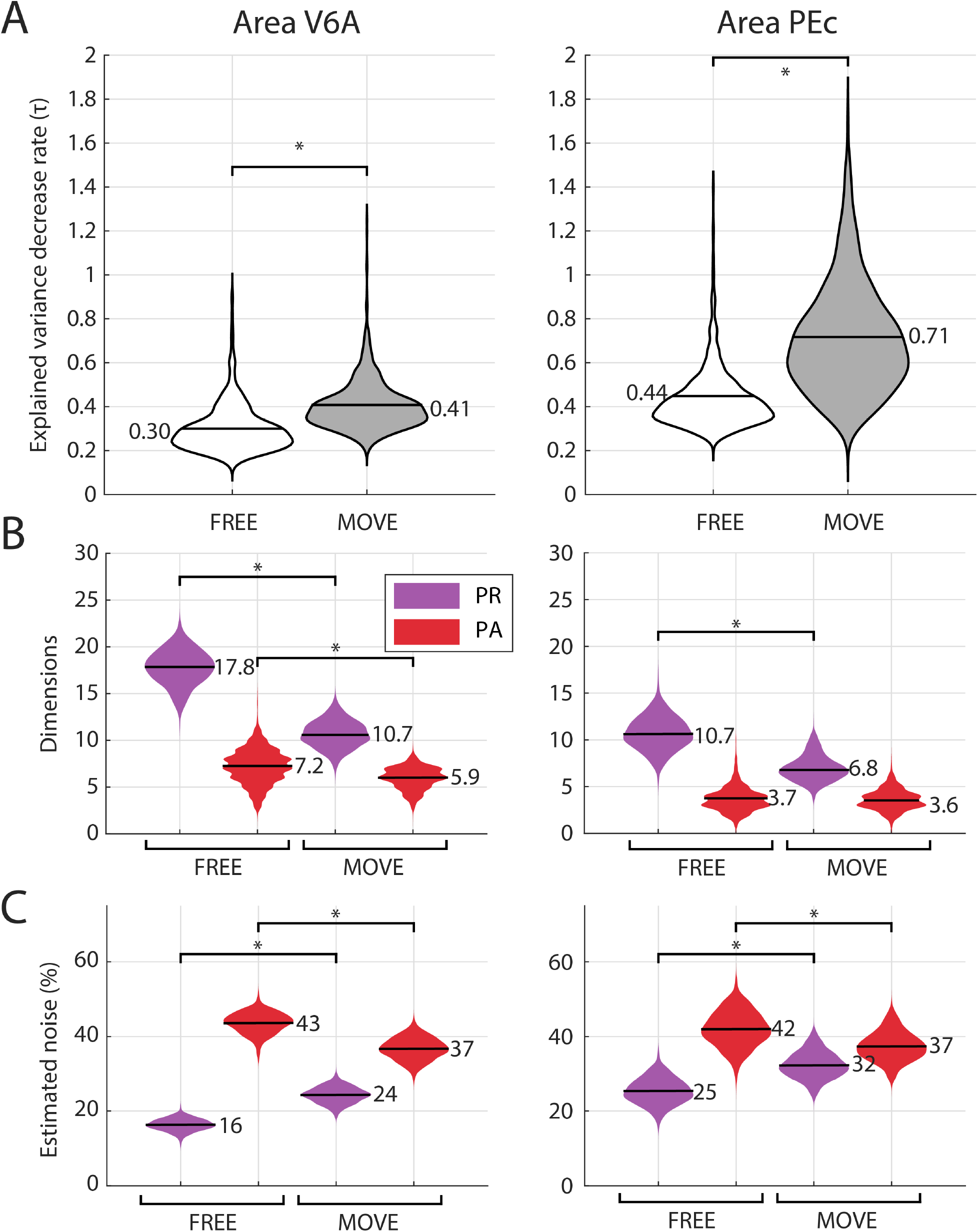
Application to real parietal activity (V6A and PEc areas). A) The decay rate (*τ*) of the variance explained by the PCs is calculated on the bootstrapped data matrices. B-C) Participation Ratio (PR) and Parallel Analysis (PA) were applied for dimensionality and noise estimation. Data spanned the first 400ms after trial beginning (FREE) and the first 400ms after movement onset release (i.e., movement onset, MOVE). Neural activity was calculated in 50ms bins, condition averaged, soft normalized and mean centered. Data were collected from Monkey F (Diomedi et al., 2024a, 2024b). Black horizontal lines in the violin plots indicate the mean value. Asterisks indicate significant differences (Wilcoxon test, p<0.01). The PA distributions in B) show irregular edges because the dimensionality estimated by PA can only assume discrete values.

Indeed, the dimensionality assessed by PR decreased from FREE to MOVE (mean values = [17.8 and 10.7] vs. [10.7 and 6.8] for V6A and PEc respectively, see Fig. 6B), with statistically significant differences (Wilcoxon test, *p <* 0.01). The mean explained variance was around 76%, in ac-cordance with previous results (Fig. 1), with a mean noise estimate by PR around 24% (i.e., 100 – explained variance), showing an increasing trend from FREE to MOVE (Wilcoxon test, *p <* 0.014;Fig. 6C). However, when the dimensionality was assessed using PA, the trend was much less marked or even absent: for V6A the dimensionality during FREE was still higher than during MOVE, but the difference was reduced to only one dimension (mean values = [7.2 vs. 5.9]; Wilcoxon test, *p <* 0.01; Fig. 6B); similarly, in PEc the dimensionality for FREE and MOVE was almost identical (3.7 vs. 3.6; Wilcoxon test, *p* = 0.8). Interestingly, the noise estimated by PA varied significantly across the two task phases, with more noise estimated during FREE ([43 and 42%] in V6A and PEc, respectively) rather than during movement ([37 and 37%]; Wilcoxon test, *p <* 0.01; Fig. 6C), in contrast to what was observed using PR. These findings are consistent with a population activity which is more structured (i.e., correlated across units) during the movement phase, leading to fewer, less noisy significant dimensions. Indeed, the variance explained in MOVE was reduced by about 7 − 8%, despite using 40% less dimensions, according to PR estimates. According to PA, the variance explained in MOVE increased by 6 − 7% with 18% less dimensions for V6A or with the same number of dimensions for PEc (Fig. 6). Note that given the matrix dimensions and the *τ* range (approx. 0.30*−* 0.70), the dimensionality estimated on the real data is in accordance with our simulations designed to identify the dimensionality upper bound. Indeed, for PA we expected 7 *−* 8 identifiable dimensions with *τ ≃* 0.3 and 4 *−* 5 with *τ ≃* 0.7, depending also on the quantity of noise (Fig. 5A). For PR, instead we expected 6 − 21 identifiable dimensions with *τ≃* 0.3 and 3 9 with *τ* 0.7, with a much stronger dependence on the noise in the data (results in the online repository related to this work). In sum, the method chosen for retaining the latent variables can significantly influence the outcome of an analysis, underlying the need to carefully justify the rationale leading to the choice of a specific criterion (see Discussion).

## Discussion

### Best methods to reliably estimate dimensionality

In this work, we investigated through extensive simulations the performance of different criteria to choose the optimal number of PCs in data with a known underlying structure, focusing on noise sensitivity and the stability of dimensionality estimates. The commonly used fixed variance thresholds (80 *−* 90%) guaranteed good results only when the noise regime was close to the predefined threshold, with performance degrading rapidly with increasing or decreasing noise. Moreover, these thresholds methods are extremely sensitive to the number of units in the dataset, often overestimating dimensionality as the unit count increases. The Participation Ratio behaved similarly to a 75 *−* 77% threshold, but offered more flexibility, mitigating some of the hard threshold limitations (e.g., it exhibited lower sensitivity to number of units / variables in the data matrix or noise). However, PR cannot provide an independent estimate of the dataset dimensional-ity from the amount of noise it contains, overestimating noise in low-noise conditions (similarly to the 80% threshold) and underestimating it in high-noise conditions. The Kaiser Rule performed quite well with long time series (i.e., sufficient rows in the data matrix, 1000+) where the impact of spurious correlations was reduced, but it was still sensitive to the amount of noise and to the number of units. Cross-validation and parallel analysis resulted to be the most reliable criteria. In particular, PA was more conservative, ensuring a robust performance across different noise regimes and data normalization methods. The cross-validation we implemented produced slightly better average estimates in low-noise regimes. Note that all of these findings were consistent for simulations designed to mimic both neural data sets and more standard data sets. PA was prone to a statistical bias as it tended to underestimate dimensionality and overestimate noise when not many simulated neurons were available, whereas with a larger number of units both the dimensionality and noise estimates converged towards the truth values (e.g., showing a good consistency). Instead, cross-validation proved to be an estimator slightly less, but it requires much more computational load. Its estimates approach the ground truth with a larger number of available units. Regarding consistency, note that this was not influenced by the length of the time series (Fig. 4), but rather by the larger number of neurons available (importantly, as long as the neurons added are always combinations of the same latent variables, as in the case of our simulations; Fig. 3).

For future applications, a combined approach using PA and cross-validation seems to be the most appropriate. The superiority of PA compared to other criteria is in accordance with previous works (Dinno, 2009; Patil et al., 2008) and has been recently confirmed through simulations of motor cortex neural data (Altan et al., 2021). However, in this latter study, the authors did not address two relevant points: different shapes of the explained variance distribution across PCs and the possibility to use retention criteria to simultaneously evaluate the noise of the data. Here, we found that the parallel analysis / cross-validation performance is influenced by the shape of the curve, with higher decay values preventing a clear distinction between informative and noise components for many PCs (see below). Both methods also provided extremely accurate noise estimates. An interesting, but quite complex alternative procedure for retaining only significant components can be found in the work of Perich et al. (2018), which is based on the estimation of trial-to-trial noise through reshuffling. This method shares similarities with PA but was not adopted in our work due to the extensive literature supporting PA and its well-established background.

To summarize, based on our results, we strongly discourage the use of fixed variance thresholds, whereas we suggest that methods such as parallel analysis and cross-validation might be widely adopted in Neuroscience.

### Upper bounds for real data applications

The choice of the correct number of dimensions can have important implications on data interpretation (e.g., when trying to infer the information content of neural dynamics that may actually be dominated by noise) and on subsequent analysis (e.g., when calculating measures such as the tangling index or the alignment between subspaces; Russo et al., 2018; Elsayed et al., 2016), thus representing a critical step. To provide guidance for future applications with real data, we designed a subset of simulations with a complex underlying structure (i.e., with a high number of latent variables) to explore scenarios where not all latent variables would be detectable in order to estimate upper bounds for dimensionality. The rationale behind this analysis is that given a known decreasing curve in the explained variance for the PCs and a data matrix with specific dimensions, one cannot reliably estimate an arbitrary large number of significant components. Here, ‘significant components’ refers to those components that account for a fraction of variance significantly higher than spurious correlations and noise. Accordingly, as the ‘slope’ *τ* of the curve increases (indicating that the variance is more concentrated in the first PCs) and the matrix dimensions decrease (raising the likelihood of random spurious correlations), the number of significant components that can be estimated decreases. We focused on results provided by PA, since it performed reasonably well across a variety of scenarios in our previous simulations (results for other matrix dimensions or other criteria can be conveniently consulted via a Web app with a simple graphic interface, see ‘Code availability’). Given a physiological range of *τ* (0.5 – 0.8;(Qi et al., 2022; Jiang et al., 2020; Kobak et al., 2016)) and assuming noise in the range 5°50%, the maximum number of dimensions that can be reliably estimated is 2 *−* 5 for small matrices and 4°6 for larger ones, even when the actual dimensionality of the simulated data is much higher (from 20 to 80 components). Since PA is quite conservative, a few more components could be considered in specific cases. The wide noise range used in our simulations, even reaching very high values (up to 50%), may seem unphysiological, but i) in real situations we do not know a priori how much noise will be present in the system, so we wanted a simulation range wide enough to provide guidance in multiple contexts; ii) according to the definition we have adopted here of ‘noise’ (i.e., uncorrelated neural activity), it is not so unlikely to reach such high noise levels, due to different tuning curves across neurons, mixed selectivity, etc. (see for example our results on real neural data); iii) since PA is almost completely unaffected by noise as far as dimensionality estimation is concerned (Fig. 1A and 5), the final results would not be particularly different in the case of a more conservative, narrower choice of noise range. According to these findings, the retention of more PCs in future studies should be carefully evaluated on a case-bycase basis, entailing the risk of considering essentially more noise than actual neural dynamics.

### Real neural data application

We applied both the PR and PA criteria to real spiking data, to estimate the neural dimensionality of parietal activity in two different phases of a reaching task (namely during FREE and MOVE). We found that the dimensionality of the population activity decreases when it is estimated by PR passing from FREE to MOVE phases, whereas it is much more stable when assessed by PA.

These apparently contradictory results can be reconciled by considering how the different methods implicitly define the dimensionality of a matrix. As already observed on simulated data, the PR behaves similarly to a variance threshold, thus indicating how many dimensions are required to account for a certain amount of variance, regardless of whether these dimensions can be distinguished by noise (i.e., uncorrelated activities) or spurious correlations. In contrast, the PA focuses on estimating how many components can be discriminated from noise, disregarding the total amount of variance explained which in turn is provided as a corollary result and allows for the estimation of noise.

One hypothesis is that stimulus presentation or motor behavior (in our case, a reaching movement) strengthens the structure of the population activity, increasing correlations across units and enhancing shared neural dynamics. In this situation, more variance is expected to be concentrated in the first dimensions. Accordingly, the ‘dimensionality’ estimated by PR (and all similar variance threshold criteria) will decrease because fewer components are required to reach the variance threshold. On the other side, the ‘dimensionality’ estimated by PA (but also by crossvalidation, for example) would remain stable (or even increase) because a higher number of components will account for enough variance to become distinguishable from noise. This hypothesis is in line with this work and previous results (Mazzucato et al., 2016), but it may be also related to the neural variability, quenching due to stimulus presentation observed at single cell level (Churchland et al., 2011, 2010).

These observations and the other results here presented highlight an ambiguity in the definition of the term ‘dimensionality’ when it applies to neural data. Indeed, ‘dimensionality’ is normally used to indicate the number of so-called ‘neural modes’ which are defined as latent, independent patterns of activity that “capture a significant fraction of population covariance” (Gallego et al., 2017). Unfortunately, the common use of variance thresholds and PR is in contrast with this definition, since these criteria do not retain components based on their significance as defined above nor on their covariance in the population, as mentioned, but simply on the total explained variance. Conversely, methods (such as PA and cross-validation) that do retain components based on their significance or on their covariance in the population seem to be in line with the current definition of dimensionality, but they are rarely (or never at all) used. Thus, the way dimensionality is estimated in practice in much of the literature seems to be in overt contradiction with the definition of ’dimensionality’ itself (and this could incidentally produce results opposite to those of the methods that seem more appropriate to the task). Given this ambiguity, the term ’dimensionality’ should be carefully defined on a case-by-case basis, while further theoretical work could aim to resolve this issue. The terms ’structural dimensionality’ or ‘dimensionality of the latent structure’ could be introduced to refer to the number of dimensions that can be uniquely identified with respect to spurious correlations or noise by methods such as PA and cross-validation. In this sense, a neural dataset could be charac-terized by an ’apparent dimensionality’ (equal to the number of observed variables, i.e. the single neurons; also named ’ambient dimensionality’ by Jazayeri and Ostojic, 2021), a lower ’latent dimensionality’ (equal to the number of latent variables identified by thresholds/PR methods and basically indicating the mere concentration of variance in the first components) and an even lower ’structural dimensionality’ (equal to the number of latent variables identified by PA/CV methods and providing hints on the real structure can be significantly discriminated in the dataset). For example, these concepts play a crucial role in defining ’embedding’ (determined by linear methods such as PCA and thus strongly influenced by the criterion used) versus the ’intrinsic’ (determined with nonlinear methods) dimensionality of neural data (Fortunato et al., 2024; Jazayeri and Ostojic, 2021). Finally, from the application of PA (which demonstrated a good ability to reliably estimate the amount of noise) on real data, we have an estimate of the amount of noise (in the range 37% – 43%) that is much higher than what might be expected. This further justifies our choice to use such a wide noise spectrum (0 *−* 50%) during the previous simulations.

### Limitations and future perspectives

In this work, we focused on PCA as the most common linear dimensionality reduction technique. This was a deliberate choice since this method is much simpler to understand and use compared to the more complicated nonlinear methods. Nonetheless, it still constitutes a powerful tool for Neuroscientists and much influential work used PCA or its derivations (Pellegrino et al., 2024; Heller and David, 2022; Keemink and Machens, 2019; Semedo et al., 2019; Lara et al., 2018; Seely et al., 2016; Kobak et al., 2016). Furthermore, PCA is the standard for determining the so-called ‘embedding dimensionality’ (Humphries, 2023; Jazayeri and Ostojic, 2021) and it is supported by a strong theoretical background (Gao et al., 2017). Similarly, we decided not to include other (possibly more robust) methods to estimate the optimal number of PCs, such as the Empirical Kaiser Criterion proposed by Braeken and van Assen (2017), for reasons of simplicity and because they are still supported by a limited literature.

Here, we tested a number of criteria that can be used in a variety of PCA applications. For example, while parallel analysis can be adapted only for all those techniques involving a covariance matrix and/or eigenvalues, the cross-validation procedure we have used can easily be extended to other types of analysis (see a similar suggestion in Cunningham and Yu, 2014). This could be the case, for example, with the Principal Component regression (PCR), the Partial Least Square (PLS) regression or the Canonical Correlation Analysis (CCA) which are gaining popularity for modelling interarea communication subspaces or the relationship between muscle and neural activity (Veuthey et al., 2020; Ames and Churchland, 2019; Ruff and Cohen, 2019; Kaufman et al., 2014). In these cases, very complex procedures are required to identify the significant latent components (see for example Veuthey et al., 2020), while the cross-validation scheme we implemented here, removing data along time and neuron dimensions simultaneously, could lead to more robust re-sults in a single step. Outside neural data analysis, but still in Neuroscience domain, the identification of muscle synergies from EMG data is mainly performed with non-negative matrix factorization (Rabbi et al., 2020) or its more efficient variants (Scano et al., 2022). However, determining the number of synergies to be retained is still problematic and the same cross-validation scheme could be beneficial.

## Conclusion

In this study, using a rich variety of simulated data, we compared different criteria that estimate the number of principal components. Hard explained variance thresholds and Participation Ratio are the most used in Neuroscience, but their results were extremely affected by the presence of noise and the number of variables (i.e., recorded neurons) in the system. Among the other criteria tested, Parallel Analysis and cross validation were the least influenced by noise and available neurons, representing promising tools to further applications to real-world data. As a clarification, our findings by suggesting the use of parallel analysis or cross-validation for estimating data dimensionality (and the amount of noise) are not intended to question any of the previous studies that have adopted other retention criteria. On the contrary, we meant to add support for the future use of more robust methods in a collaborative direction and possibly open new perspectives on how to interpret neural data. Together with this manuscript, we also share software to facilitate the adoption of the criteria presented here and to provide tools that can be used routinely as part of preliminary analyses.

## Data availability

Real neural data recorded from V6A-PEc parietal areas are available at the G-node data sharing website at https://doi.gin.g-node.org/10.12751/g-node.7q2dbp/ (Diomedi et al., 2024b). For an extensive description of the datasets see Diomedi et al., 2024a.

## Code availability

The MATLAB software package for applying various dimensionality estimation criteria can be found at https://github.com/francescovaccari/Dimensionality-estimation. All the results of simulations designed to provide upper bounds on dimensionality according to different criteria (some of which were used to generate Fig. 5) can be found at https://github.com/francescovaccari/Dimensionality-estimation-upper-bound.

## ACKNOWLEDGEMENTS

Thanks to the Machens Lab team for the fruitful feedback.

