## Supplementary Materials for "More or fewer latent variables in the high-dimensional data space? That is the question"

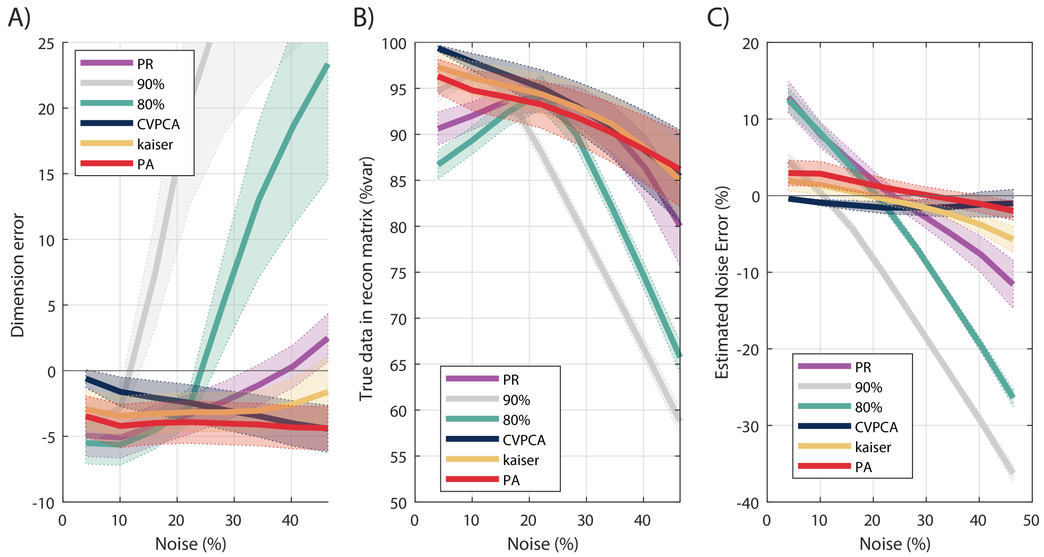


**Fig. S1.** Performance of different criteria for selecting the optimal number of Principal Component tested on simulated data (RANDOM SOURCE SET) as a function of noise (simulation parameters were set according to Table 1). The metrics calculated were: the dimension error, i.e. the difference between the estimated number of PCs and the number of latent variables $N$ used to generate the data (A); the information about the latent variables retained in the denoised matrix (B); the estimated noise error (C). Other conventions such as in Fig. 1

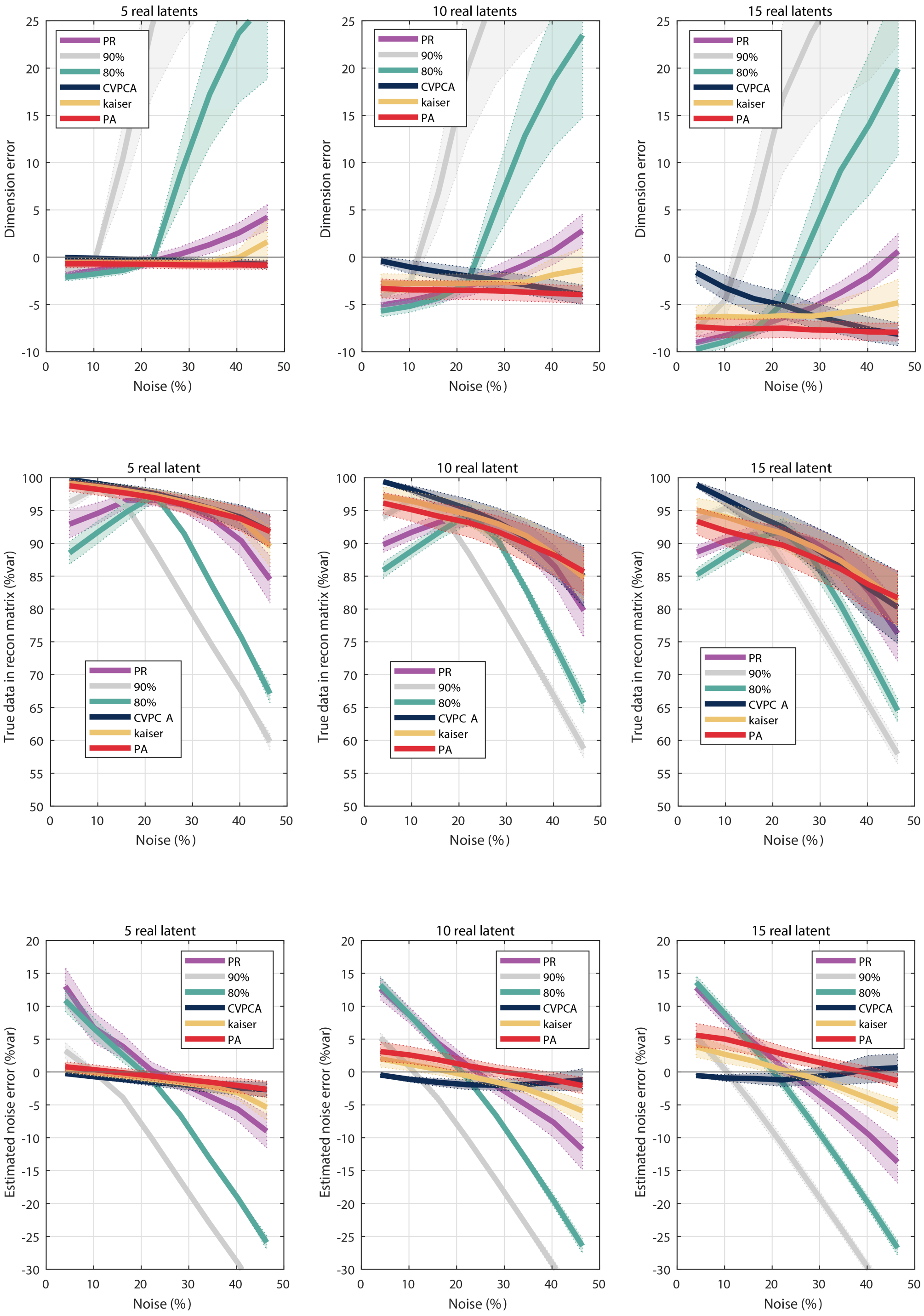


**Fig. S2.** Performance of different criteria tested on simulated data as a function of noise. In this case, results are reported grouping simulations according to the number of latent variables used to generate the data matrices ($N = 5, 10$ or $15$). Other simulation parameters were set according to Table 1 (RANDOM SOURCE SET). Other conventions as in Fig. 1 and Fig. 2.

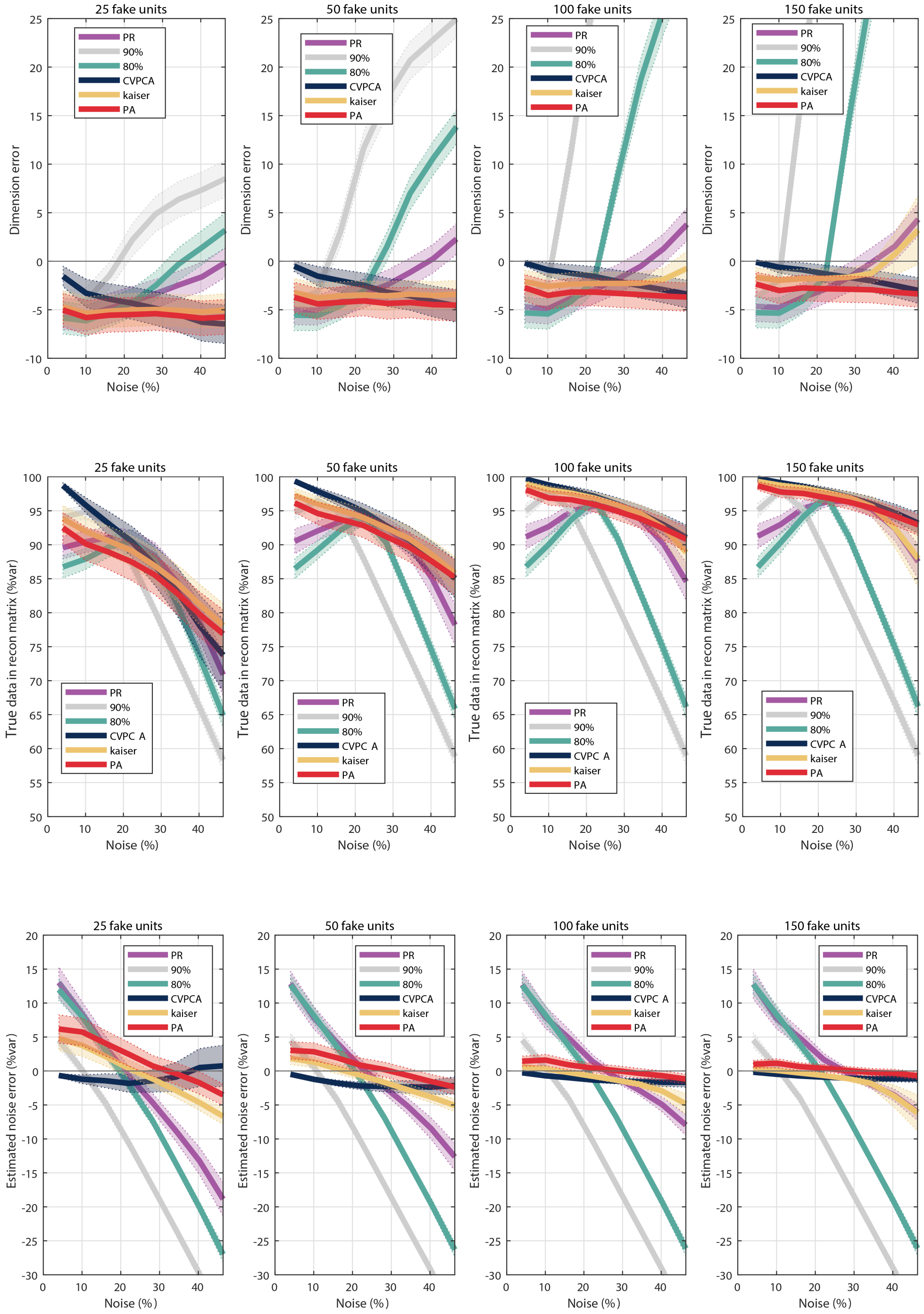


**Fig. S3.** Performance of different criteria tested on simulated data as a function of noise. In this case, results are reported grouping simulations according to the number of synthetic units ($M = 25, 50, 100$ or $150$). Other simulation parameters were set according to Table 1 (RANDOM SOURCE SET). Other conventions as in Fig. 1 and Fig. 3.

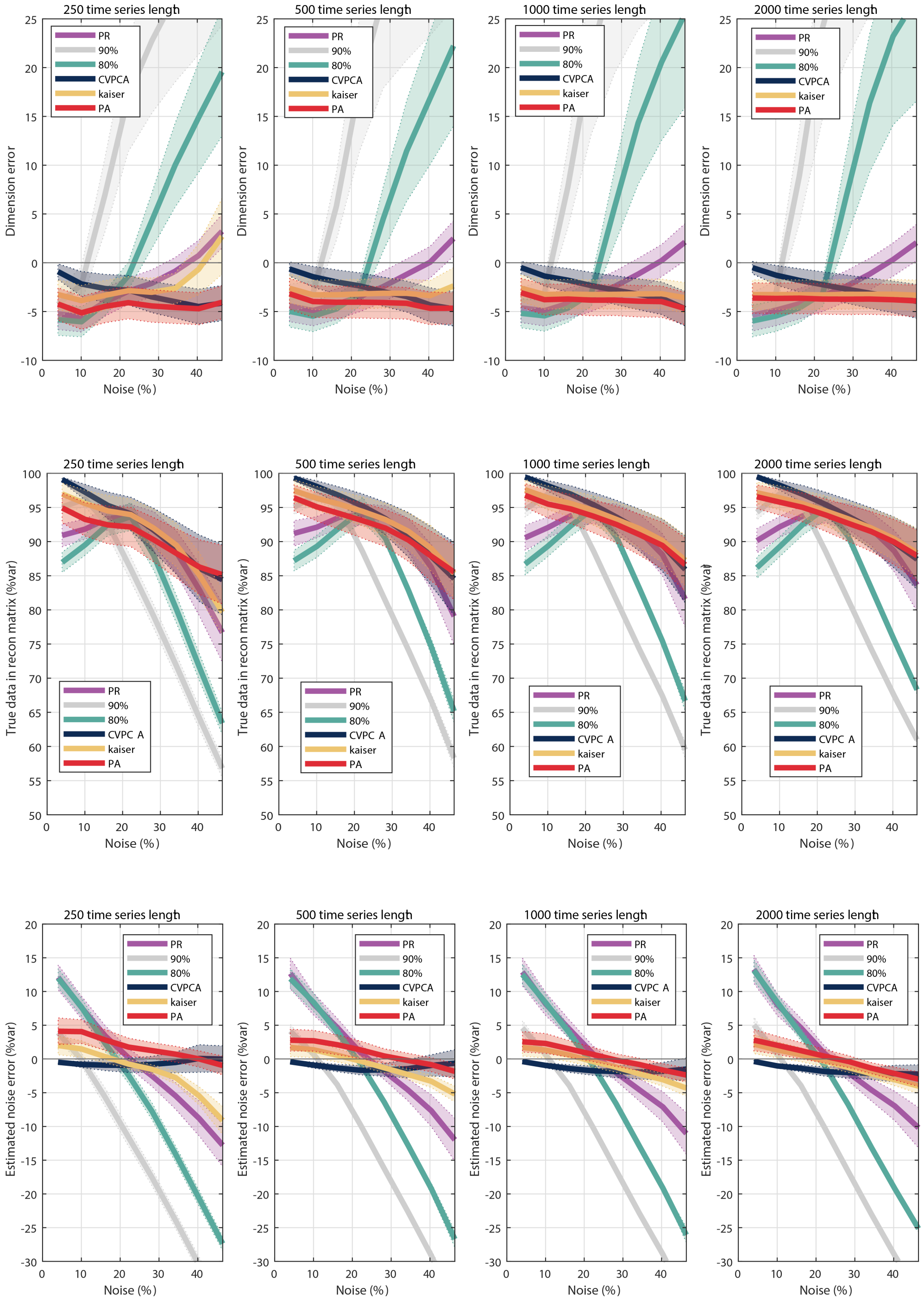


**Fig. S4.** Performance of different criteria tested on simulated data as a function of noise. In this case, results are reported grouping simulations according to the length of the time series ($T = 250, 500, 1000$ or $2000$). Other simulation parameters were set according to Table 1 (RANDOM SOURCE SET). Other conventions as in Fig. 1 and Fig. 4.
